# A consensus score to combine inferences from multiple centres

**DOI:** 10.1101/2022.11.10.515944

**Authors:** Hamed Haselimashhadi, Kolawole Babalola, Robert Wilson, Tudor Groza, Violeta Muñoz-Fuentes

## Abstract

Experiments in which data are collected by multiple independent resources, including multicentre data, different laboratories within the same centre or with different operators, are challenging in design, data collection and interpretation. Indeed, inconsistent results across the resources are possible. In this paper, we propose a statistical solution for the problem of multi-resource consensus inferences when statistical results from different resources show variation in magnitude, directionality, and significance. Our proposed method allows combining the corrected p-values, effect sizes and the total number of centres into a global consensus score. We apply this method to obtain a consensus score for data collected by the International Mouse Phenotyping Consortium (IMPC) across 11 centres. We show the application of this method to detect sexual dimorphism in haematological data and discuss the suitability of the methodology.

## Introduction

Measuring response to a treatment based on data collected from multiple resources, such as multicentre clinical trials or animal experiments, benefits from (1) lower noise level, because results are not strongly resource-dependent [1], and (2) effectiveness, because they apply to a broader population [2,3]. In these experiments, obtaining a global consensus in the statistical inference across resources is desired. However, even in highly controlled experiments, it is not always possible to control for all sources of variation across all resources. This makes aggregating statistical results from multiple resources challenging because the results may be vulnerable to biases, which lead to inconsistent inferences. The design of the study, sample size, power of the analysis, variation across centres or over time [4] and unknown errors are examples of factors that pose a challenge to obtaining a global statistical conclusion across resources [5–7]. Other confounders are the equipment that is used to perform the measurements in different resources (e.g., centres, laboratories, etc.), the level of experience of the staff and more complex environmental factors that typically arise in animal tests, such as diet, litter, handling, circadian rhythm, housing and husbandry. Therefore, in multi-resource experiments, it is crucial to control for as many variables as possible, to be able to reach global agreements [4,5,8,9]. Table 1 shows some examples of possible outcomes when an experiment is conducted in 4 centres.

**Table 1.** Examples of possible outcomes when a global inference from statistical results obtained from multiple centres is desired. In this table, we focus on the treatment effect size and p-values from centres and assume that the experiment is highly controlled and conducted by 4 centres (e.g. laboratories).

| Scenario | Setup | Inference |
| --- | --- | --- |
| 1 | All centres achieve significant results | Global consensus |
| 2 | 2 centres achieve significant results | Not clear |
|  | 2 centres did not achieve significant results |  |
| 3 | 2 centres achieve significant results | Not clear |
|  | 2 centres did not achieve significant results but 1 of them has a borderline p-value |  |
| 4 | 2 centres strongly achieved significant results | Not clear |
|  | 2 centres did not achieve significant results with p-values strongly diverting from the significant level |  |
| 5 | All centres achieved significant results but 2 in the positive and 2 in the opposite direction | Not clear |
| 6 | 2 centres achieved significant results | Not clear |
|  | 1 centre achieved significant results in an opposite direction |  |
|  | 1 centre did not achieve/borderline significant results |  |

In this paper, we present a methodological approach which seeks to find a solution to the problem of multi-resource consensus with a focus on multicentre experiments. The proposed method allows calculating a global consensus score for the effect of interest (i.e., research questions, e.g., genotype, sexual dimorphism, bodyweight effect) in multicentre studies. The method takes into consideration the number of centres where the test of interest is performed at, the direction and magnitude of the effect size and the significance level obtained from individual centres and combines the values into a global consensus score. We apply our method to data obtained by the International Mouse Phenotyping Consortium (IMPC), a transnational multicentre endeavour that screens the phenotypes of single-gene knockout mouse lines and wild-type mice to understand gene function [10].

## Method

There are several approaches typically used to aggregate inferences from multicentre data. Among them, three major methods involve adjusting for centres using fixed and random models; or analysing each centre separately and then combining the results using meta-analyses [2,11–14]. Other methods are utilising group decision-making processes, such as the DELPHI method [15,16]; or using a simple majority rule criteria, such as *all centres agree* versus *at least one centre disagree;* or employing simple statistics or probabilistic criteria, such as *more than half/mean/median centres/results agree* or simple statistical tests such as T-test or ANOVA [17]. Latter approaches may suffer from insufficient power, individual bias (such as misjudgements or making decisions based on insufficient information) and may have strong underlying assumptions as well as require a large M, the total number of centres, to converge to the true inference [2,18].

Here we propose an alternative approach which combines the corrected p-values (q-values), which we obtained using the FDR [19–21], and the effect sizes from individual centres and compares them with a set of expected values as below:

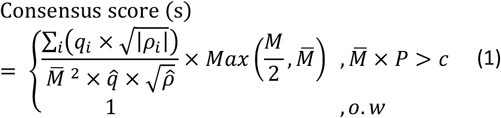

where *i* = 1, 2,…,*M* represents the *i^th^* centre from a total of *M* centres, 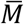 the total number of centres where the test is performed at (*M* is not necessarily equivalent to 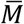 in multicentre multi-test studies where the aim is to compare several measurements across centres while fixing the number of centres), *q_i_* the corrected p-value (q-value) from the statistical test performed in centre *i* for the effect of interest (e.g. sex, genotype, body weight effect, etc.), *ρ_i_* the estimated standardised effect size from the statistical test that is performed in centre *i*, such as Cohen’s *d* effect size [22] and 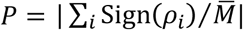 is a penalty term to control for the directionality of the results, and the Sign(*ρ*) is the sign function defined by

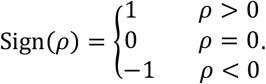

Finally, *c*, 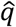 and 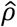 are the minimum required number of centres for the analysis, the expected q-value and effect size from the prior information, respectively. We recommend *c* = 3, 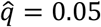 and moderate expected effect size 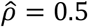 [3,23,24] as the preliminary values for high-throughput experiments, such as in the IMPC. Figure 1 shows the distribution of the standardised effect sizes for the IMPC haematological traits and empirical mean (10% trimmed) from the data and the recommended expected effect size, 0.5. We stress that the choice of these values should be based on prior information. We further assume that (1) there is no unusual temporal variation in the data (Supplementary Fig.1), (2) the statistical tests are consistent and sufficiently powerful and adequate for the data under study, (3) the method to adjust the p-values is adequate (e.g. FDR); and (4) the effect sizes are estimated from the normalised data. Here normalising data refers to performing the statistical analysis on the standardised data as below:

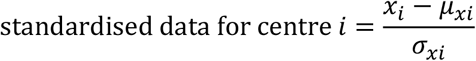

**Figure 1.**
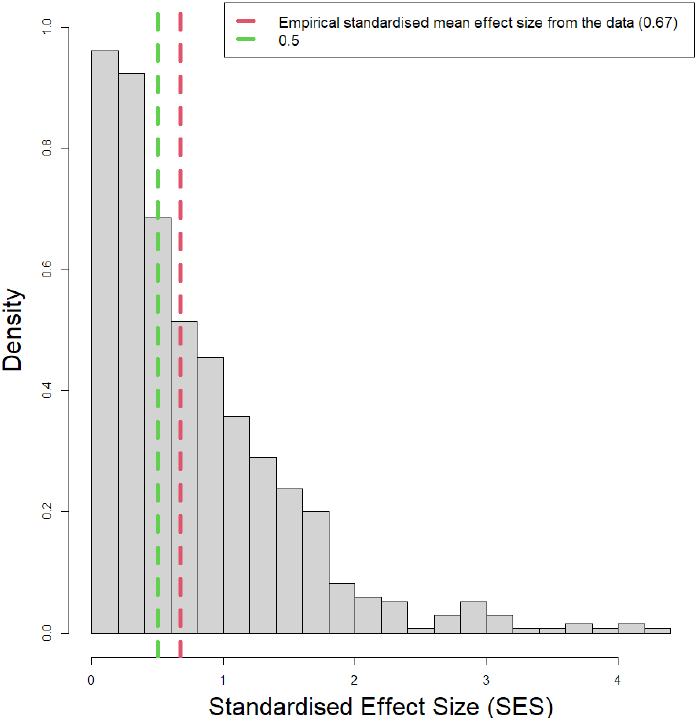
The distribution of the standardised effect sizes (SES) for the IMPC haematological traits. The empirical 10% trimmed mean SES (dashed red line) is 0.67 and the recommended value for the expected effect size 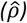 is 0.5 (dash green line).

Where *x_i_*, *μ_xi_* and *σ_xi_* are the raw values, mean and standard deviation of the data from centre *i* respectively. The resulting scores from Eq.1 range in the (0,+∞) interval and the agreement of the multicentre statistical results can be evaluated by using −log(s) so that

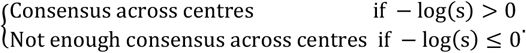

The magnitude of −log(s) from Eq.1 is not bounded. As a result, a larger value in the positive (or negative) direction reflects a stronger agreement (or lack of agreement) among resources. For the special case where −distribution of the standardised effectlog(s) = 0, one can conclude that either there is not enough information in the data to calculate the scores or there is not enough agreement across centres. Throughout this paper, we use the term “not enough agreement” in contrast to “disagreement” to emphasize the difference between strong detection of consensus and not finding enough evidence to establish consensus among centres. Table 2 shows several scenarios as well as the inferences from the scores in Eq.1. This table shows that the most ambiguous scenario happened when all centres achieved the same effect size and q-value to the expected values (scenario 2) or the centre achieved a range of opposite (in sign) effects so that *M* × *P* ≤ 3 (scenario 3). Because q_j_ and p_j_ are continuous real values, q_j_, |p_j_| ∈ [0, ∞), scenario 3 happens with an extremely low chance that can be safely neglected.

**Table 2.**
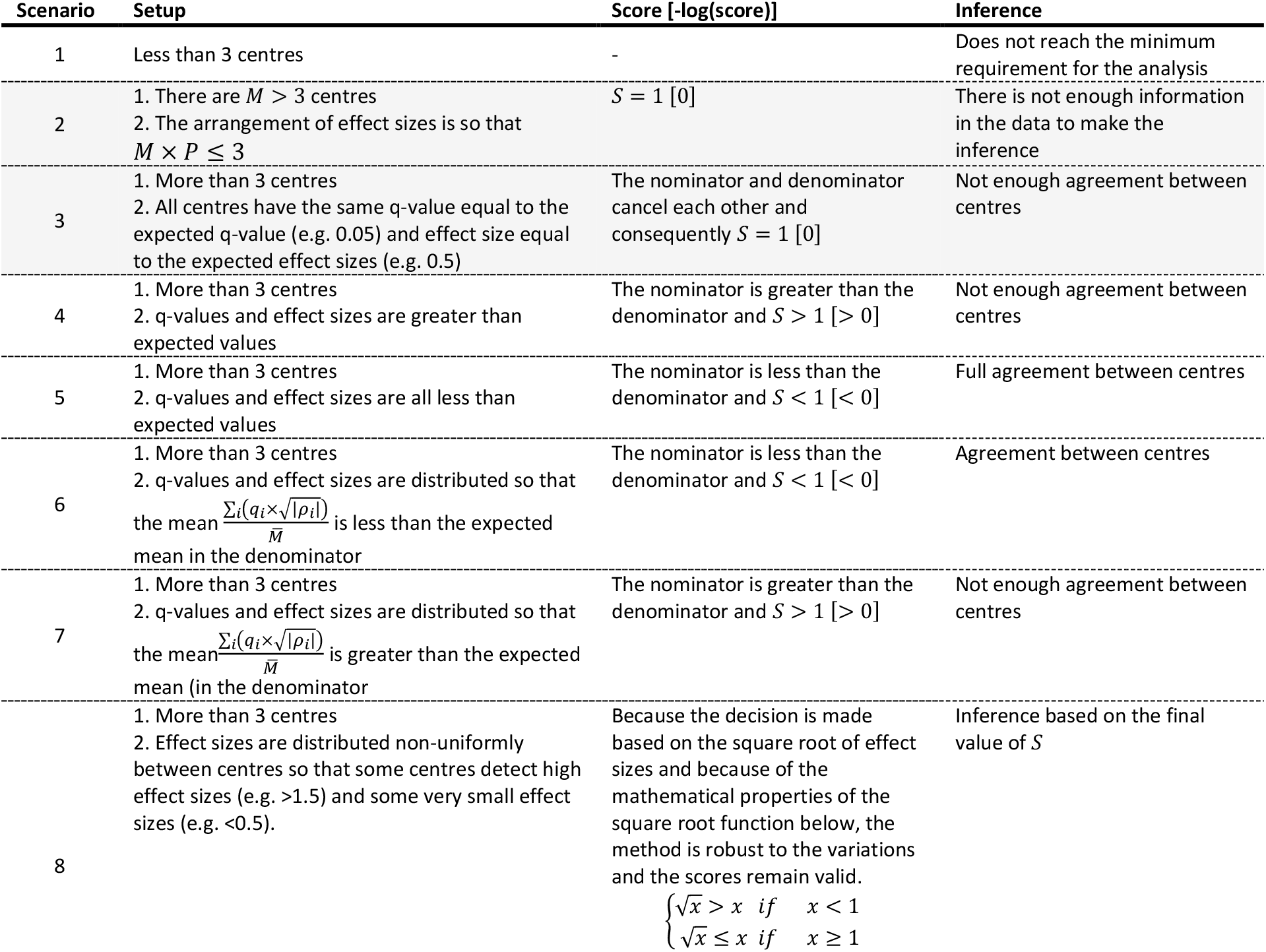
The demonstration of the scores calculated from Eq.1 in a set of scenarios with 3 or more centres when the proposed scoring method in Eq.1 leads to different values and inferences. Scenarios 2 and 3 lead directly to a score of 1 (log(s) = 0) with two different inferences: (i) there is not enough information in the data to make the inference; or (ii) not enough agreement between centres. Because q_i_ and p_i_, are continuous real values, q_i_, |p_i_| ∈ [0, ∞), In practice, scenario 3 happens with an extremely low chance and can be safely ignored. The first scenario should be detected in a pre-processing step.

## Results

In this section, we show the application of the proposed scoring method along with two methods from the literature, precisely global consensus and metadata analysis, to identify sexual dimorphism in the IMPC haematological data collected from wild-type (WT) mice, with an average age of 16-18 weeks, over a 3-year period from 1^st^ January 2018 to 31^st^ December 2020, with a minimum required threshold of 50 mice per sex. Our choice of data is inspired by the importance of the haematology parameters reflecting overall health. The data used in this study can be accessed via the IMPC web portal under the URL www.mousephenotype.org (data release 15.1 – October 2021).

The IMPC is a global effort aiming to generate and characterise knockout mouse lines for every protein-coding gene in mice [25–28]. The IMPC data are collected from several independent centres worldwide [10]. Every centre contributes to the data collection by adhering to a set of standardised phenotype assays defined in the International Mouse Phenotyping Resource of Standardised Screens (IMPReSS - www.mousephenotype.org/impress). Although all centres follow the same Standard Operating Procedures (SOPs), there may be unavoidable or necessary variations in the implementation of the experiments (such as mouse age or time of the day when the test is performed), equipment (such as manufacture, model and kits) as well as the level of expertise and experience of staff (experimenter effect), in addition to variations in inbred mouse strain (Table 3) [29]. This may lead to differing results across centres, which makes a global inference from the results challenging.

**Table 3.**
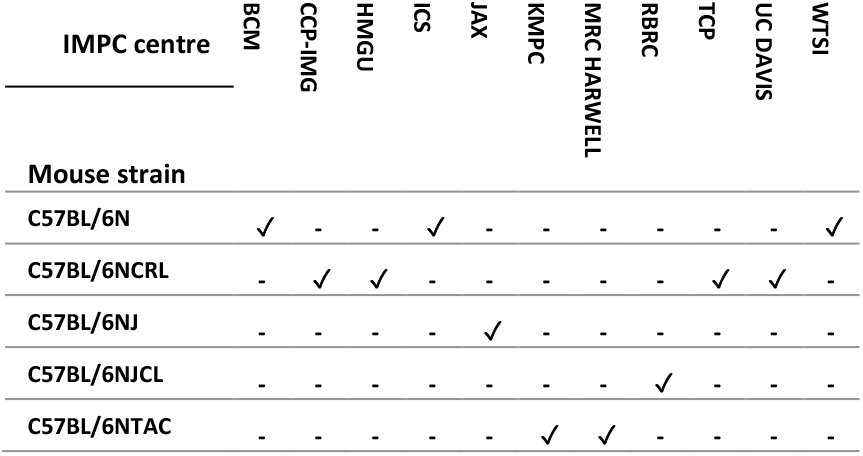
Mouse strains that are used by the IMPC centres for the haematological data collected from 1^st^ January 2018 to 31^st^ December 2020.

### IMPC haematology

The IMPC haematology procedure encapsulates 22 measurements of blood properties such as counts and concentrations (white blood cell count, red blood cell count, haemoglobin concentration, platelet counts, etc.), as well as additional and derived haematological parameters (haematocrit, mean red blood cell volume, mean red blood cell haemoglobin, mean red blood cell haemoglobin concentration, etc.). Figure 2 (top) shows red blood cell counts, (middle) the haemoglobin concentration and (bottom) the monocyte cell counts collected by IMPC centres. The shifts in the means are most likely due to differences in the equipment used to take the measurements and can be removed by normalising the data. The top plot shows consistently higher red blood cell counts in males than females across centres, whereas there is not a clear pattern for the haemoglobin concentration. For the monocyte counts, males present consistently higher values, except for one centre, which shows the opposite.

**Figure 2.**
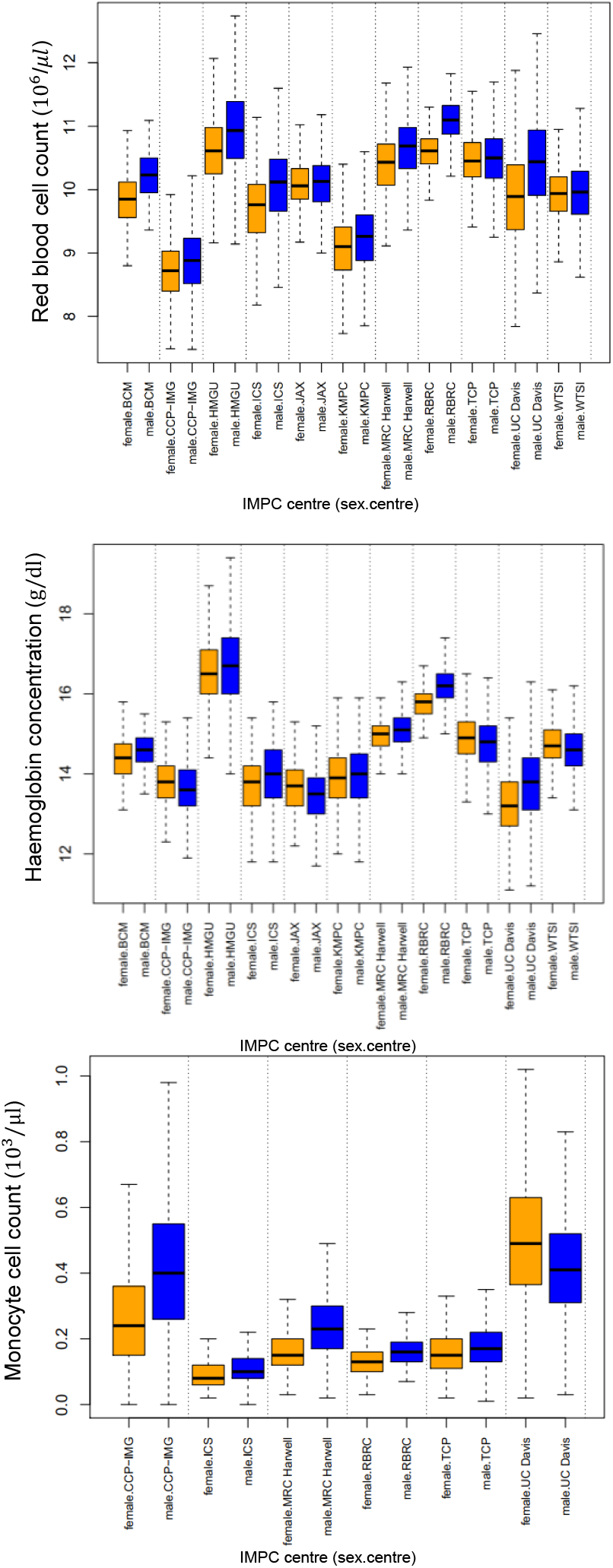
The distribution of red blood cell counts (top), the haemoglobin concentration (middle) and monocyte cell counts (bottom) for wild-type mice from the IMPC, split by sex and phenotyping centre. The orange and blue represent females and males, respectively. The consensus score for the red blood and monocyte cell count traits are respectively −log(s) = 0.30 and 2.28 which implies a global agreement across IMPC centres in identifying sexual dimorphism; the sign of the average effect size indicates whether males (positive) or females (negative) present higher values (males in this case, see Table 2). In contrast, the consensus score for the haemoglobin concentration trait is −log(s) = 0, which implies lack of agreement among the IMPC centres to detect sexual dimorphism for this parameter.

### Consensus score

In line with [3], the sexual dimorphism effect is tested for all 22 haematology traits, independently for WT mice from individual centres, corresponding to the same mouse strain and metadata group split. We used a linear mixed model described in [30,31] and implemented in the software R [32] and packages OpenStats [33]. As in [3], *Sex* and *Body Weight* in the fixed effect terms

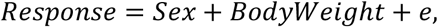

and Batch (the date when the test is performed on mice) in the random effect term. We then apply the scoring method to obtain a consensus global inference from the multicentre results, following the logic described in the flowchart below (Fig.3). We further compare our method with the global consensus criteria (all centres agree vs at least one centre disagree) and the metadata analysis proposed in [14] and implemented in the R package *metafor* [34].

**Figure 3.**
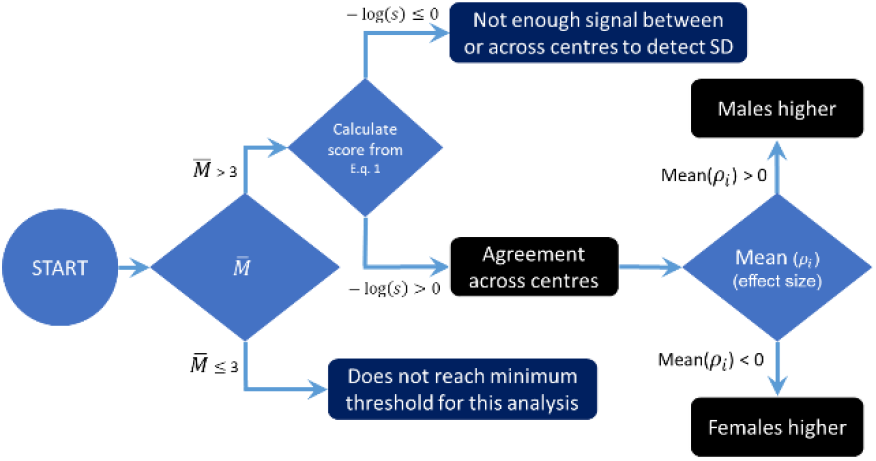
Flowchart showing the logic behind the scoring method to obtain a consensus global inference from multicentre results. The first step involves examining the number of centres performing the test; when there are more than 3 centres, the consensus score is calculated. Provided −log(s) > 0, a multicentre consensus signal is established (accepted) and the direction of sexual dimorphism based on the sign of the average effect sizes is reported.

Table 4 shows the outcome of the scoring method for the 22 haematological parameters measured by the IMPC, as well as the comparison with a consensus method based on all centres agreeing on a significant sex effect and the meta-analysis method. Using the method proposed here, there is consensus among 11 IMPC centres for 14 traits with −log(s) > 0, with males on average higher than females for 9 traits (red blood cell count, red blood cell distribution width, haematocrit, platelet count, white blood cell count, lymphocyte cell count, neutrophil cell count, monocyte cell count, eosinophil cell count) and females on average higher than males for 5 traits (mean cell volume, mean corpuscular haemoglobin, mean cell haemoglobin concentration, mean platelet volume, and lymphocyte differential count). For 8 traits, the scoring method leads to zero or negative values, reflecting a lack of consensus (6 traits), or does not reach the minimum threshold of three centres providing measurements for the results to be processed (lack of information in the data - 2 traits). The meta-analysis method shows consistent results with the scoring method, however, does not obtain the homogeneity of the statistical results across the centres for the monocyte cell count (also shown in Fig.2 bottom), lymphocyte differential count and a borderline p-value for the eosinophil cell count (p-value = 0.069) and the neutrophil differential count (p-value = 0.048). Visual inspection of the data shows that the meta-analysis has a better performance for identifying the lack of agreement in *lymphocyte differential count* whereas the scoring method outperforms this method for the *monocyte cell count*. In contrast with the two methods above, the global consensus method shows the agreement across centres for the n*eutrophil cell count and Large Unstained Cell (LUC) count where the latter does not reach the requirement of a minimum of 3 centres*.

**Table 4.**
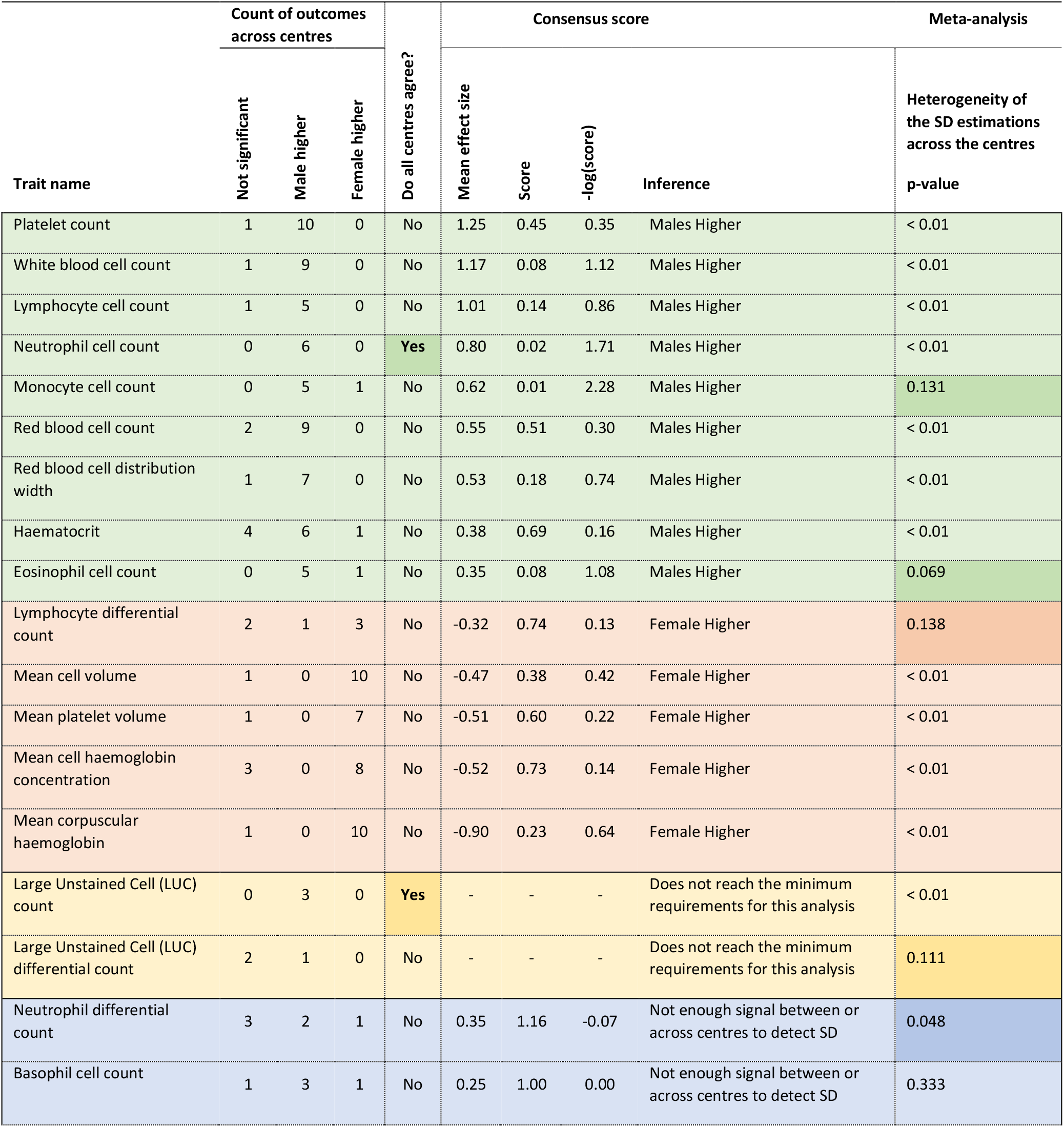
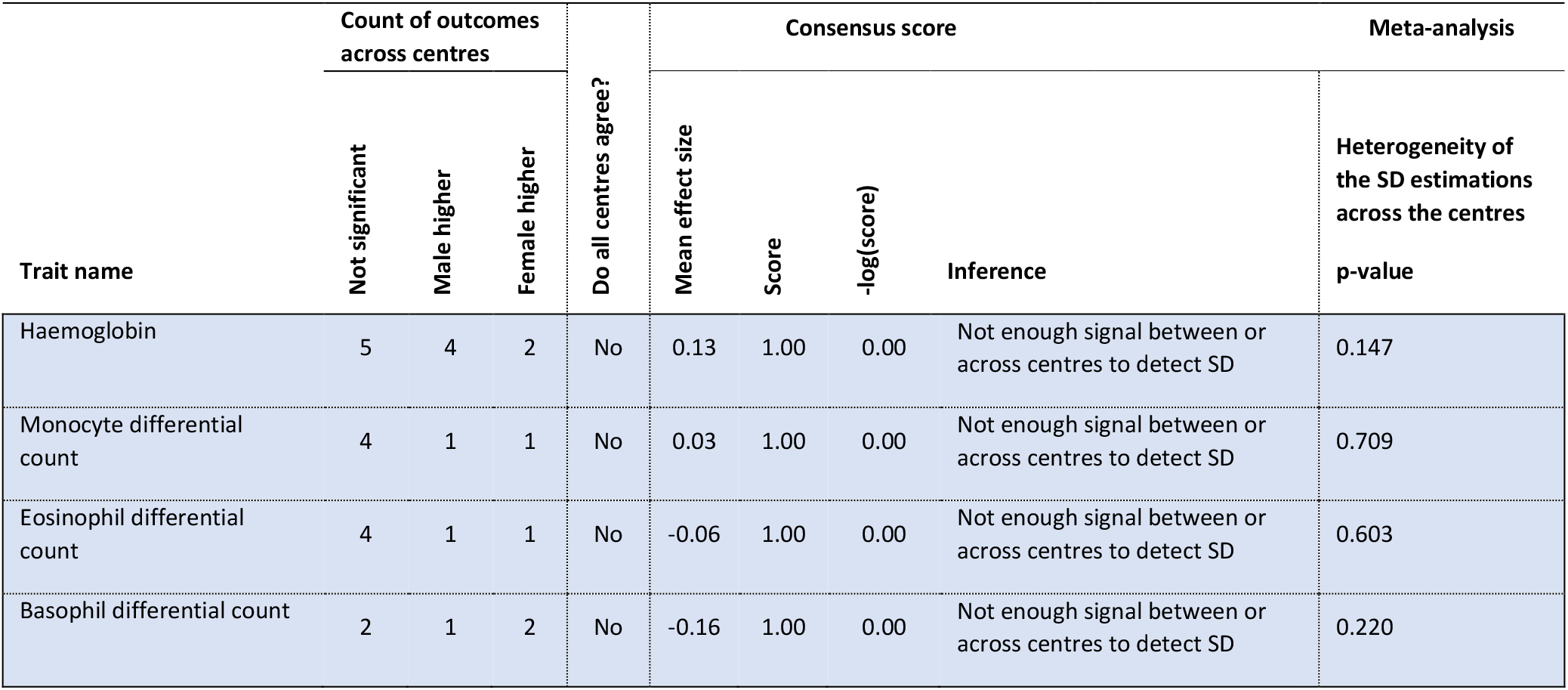
The outcome of applying the scoring method to 22 haematological measurements collected by 11 IMPC centres compared with a method based on measuring the heterogeneity of the estimates across the centres. The traits are shown in rows followed by the counts for the centre-based statistical test results, the mean effect size for the 11 centres, the consensus score and the inference, which is based on the −log(score) and the sign of the mean effect size. The scoring method identifies consensus in sexual dimorphism across centres for 14 traits (green and red rows), no agreement for 8 traits (blue rows) and 2 traits which do not meet the minimum requirements for the calculation of the score (yellow rows). Only in 2 cases, all centres agree (in bold).

## Conclusion and future work

Collecting data from multiple resources such as, in the case of this study, mouse phenotyping centres, benefits from a higher signal-to-noise ratio and a broader representation of a population. However, extra attention is required in the design and implementation of the experiments and statistical analysis to be able to make a global consensus inference from the aggregated results from individual resources [2–9,11–14,34–36]. Due to unavoidable, uncontrolled and unobserved factors, the results from all resources may only partially agree and a metric of consensus is required. In this paper, we propose a novel method which combines several aspects of multicentre experiment results including the corrected p-values, the magnitude and direction of effect sizes and the number of centres into one global consensus score.

We applied this method to identify sexual dimorphism in 22 haematological measurements collected from wildtype mice in 11 globally distributed centres forming part of the International Mouse Phenotyping Consortium (IMPC). We compared the results of this method to those obtained by the meta-analysis as well as by applying a binary method based on the agreement of all centres on the detection of sexual dimorphism. While the binary method found 2 traits reaching consensus across all IMPC centres, the method presented here allows to conclude sexual dimorphism in 14 traits, with males on average higher than females for 9 traits and females on average higher than males for 5 traits. Further, comparing our method with the meta-analysis method shows a high degree of overlap between the two 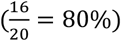 for the haematological traits. Our method shows better performance for monocyte cell count (–log(score) = 2.28 versus meta-analysis p-value = 0.131) and eosinophil cell count (–log(score) = 1.08 versus meta-analysis p-value= 0.069). However, a challenging case for the interpretation of the results is presented in comparing the outcome of the scoring method versus the meta-analysis method for lymphocyte differential count (-log(score) = 0.13 versus meta-analysis p-value= 0.138). This study has focused on the IMPC haematology traits, but we believe the approach could be applied more generally and would be suitable to assess other IMPC parameters in the future.

## Future studies

In this study, we showed the application of our scoring method to IMPC haematological data. In future studies, we will investigate the performance of the method when applied to other IMPC procedures as well as obtaining the statistical properties of the test statistic. This will allow assigning a probability of consensus to the scores (in particular when they are close to 1 or −log(score) is close to zero) that contributes to the reliability of the method.

## Declarations

## Acknowledgements and funding

We thank Helen Parkinson for her feedback on this manuscript. The research reported in this publication was supported by the European Molecular Biology Laboratory (EMBL-EBI) core funding and the National Human Genome Research Institute of the National Institutes of Health under Award Number 2UM1HG006370-11. The content is solely the responsibility of the authors and does not necessarily represent the official views of the National Institutes of Health.

## Authors’ contributions

H.H. and V.M. contributed to the development of the concept and writing of the manuscript. H.H., V.M. and K.B. contributed to the validation of the method. All authors contributed to the review of and approved the final version of the manuscript.

## Data availability

All data used in the study are publicly available via the IMPC web portal under the URL www.mousephenotype.org

---

**Supplementary Figure 1.**
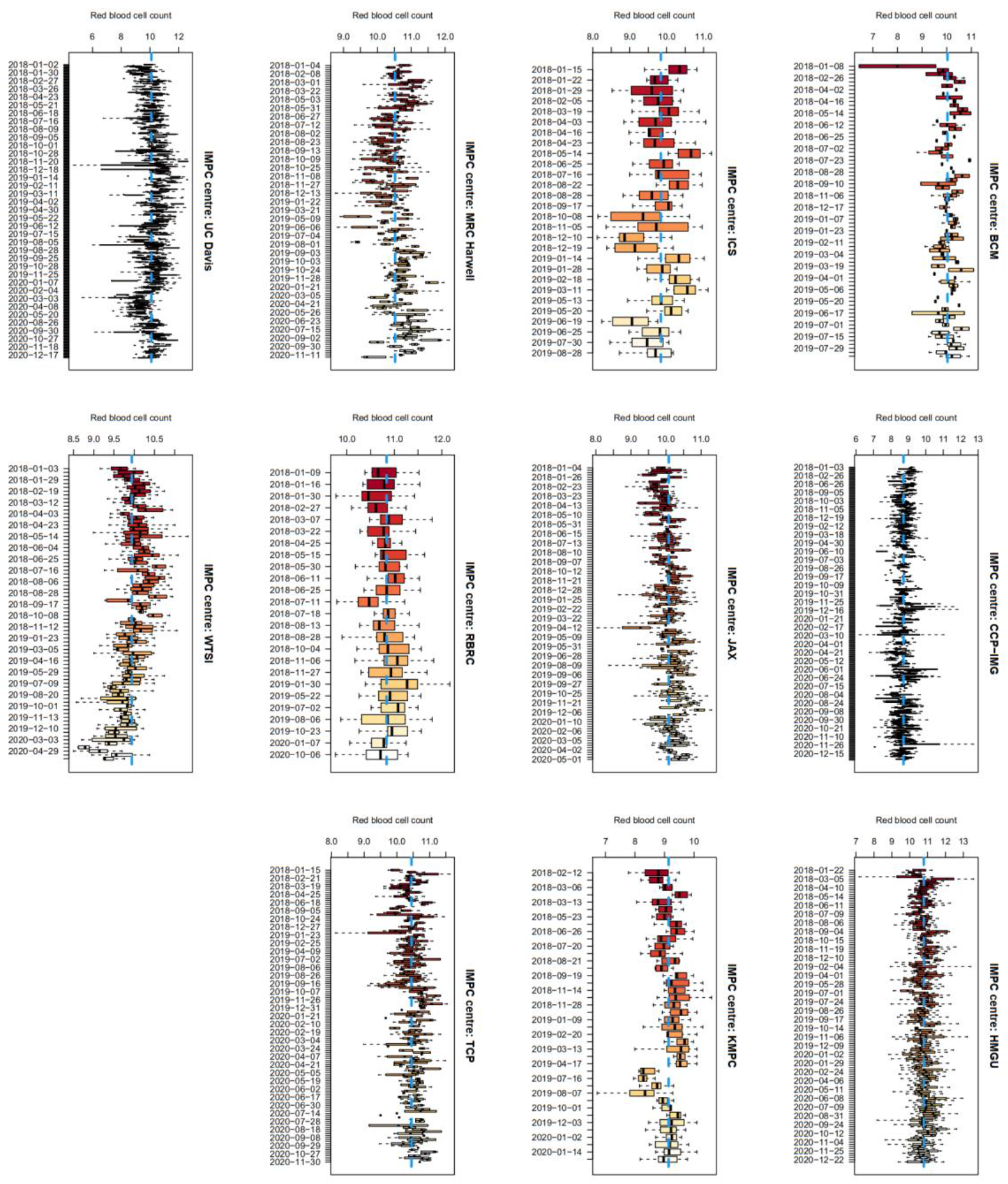
The data variation for the IMPC wildtype mice over time (batch) for the red blood cell counts across the IMPC centres. The global mean is shown by the vertical blue lines. The IMPC centres show some variations over time. Align with [1] a batch effect term is included in the analysis to account for the day to day variation in the data.

## References

1. Karp NA, Speak AO, White JK, Adams DJ, de Angelis MH, Hérault Y, et al. Impact of temporal variation on design and analysis of mouse knockout phenotyping studies. PLoS One. 2014;9. doi:10.1371/JOURNAL.PONE.0111239

2. Rashid MM, McKean JW, Kloke JD. R Estimates and Associated Inferences for Mixed Models With Covariates in a Multicenter Clinical Trial. http://dx.doi.org/101080/194663152011636293. 2012;4: 37–49. doi:10.1080/19466315.2011.636293

3. Karp NA, Mason J, Beaudet AL, Benjamini Y, Bower L, Braun RE, et al. Prevalence of sexual dimorphism in mammalian phenotypic traits. Nat Commun. 2017;8: 15475. doi:10.1038/ncomms15475

4. Haselimashhadi H, Mason JC, Munoz-Fuentes V, López-Gómez F, Babalola K, Acar EF, et al. Soft windowing application to improve analysis of high-throughput phenotyping data. Bioinformatics. 2020;36: 1492–1500. doi:10.1093/bioinformatics/btz744

5. Chung KC, Song JW, group W study. A Guide on Organizing a Multicenter Clinical Trial: the WRIST study group. Plast Reconstr Surg. 2010;126: 515. doi:10.1097/PRS.0B013E3181DF64FA

6. Hu M, Shi X, Song PX-K. Collaborative causal inference with a distributed data-sharing management. 2022 [cited 14 Oct 2022]. doi:10.48550/arxiv.2204.00857

7. Knatterud GL, Rockhold FW, George SL, Barton FB, Davis CE, Fairweather WR, et al. Guidelines for Quality Assurance in Multicenter Trials: A Position Paper. Control Clin Trials. 1998;19: 477–493. doi:10.1016/S0197-2456(98)00033-6

8. Chalmers I, Clarke M. Commentary: the 1944 patulin trial: the first properly controlled multicentre trial conducted under the aegis of the British Medical Research Council. Int J Epidemiol. 2004;33: 253–260. doi:10.1093/IJE/DYH162

9. Hogg RJ. Trials and tribulations of multicenter studies. Lessons learned from the experiences of the Southwest Pediatric Nephrology Study Group (SPNSG). Pediatr Nephrol. 1991;5: 348–351. doi:10.1007/BF00867501

10. Koscielny G, Yaikhom G, Iyer V, Meehan TF, Morgan H, Atienza-Herrero J, et al. The International Mouse Phenotyping Consortium Web Portal, a unified point of access for knockout mice and related phenotyping data. Nucleic Acids Res. 2014;42. doi:10.1093/nar/gkt977

11. Basagaña X, Pedersen M, Barrera-Gómez J, Gehring U, Giorgis-Allemand L, Hoek G, et al. Analysis of multicentre epidemiological studies: contrasting fixed or random effects modelling and meta-analysis. Int J Epidemiol. 2018;47: 1343–1354. doi:10.1093/IJE/DYY117

12. Burke DL, Ensor J, Riley RD. Meta-analysis using individual participant data: one-stage and two-stage approaches, and why they may differ. Stat Med. 2017;36: 855–875. doi:10.1002/SIM.7141

13. Bowden J, Tierney JF, Simmonds M, Copas AJ, Higgins JP. Individual patient data meta-analysis of time-to-event outcomes: one-stage versus two-stage approaches for estimating the hazard ratio under a random effects model. Res Synth Methods. 2011;2: 150–162. doi:10.1002/JRSM.45

14. Stewart GB, Altman DG, Askie LM, Duley L, Simmonds MC, Stewart LA. Statistical Analysis of Individual Participant Data Meta-Analyses: A Comparison of Methods and Recommendations for Practice. PLoS One. 2012;7: e46042. doi:10.1371/JOURNAL.PONE.0046042

15. Ven AH van de, Delbecq AL. The Effectiveness of Nominal, Delphi, and Interacting Group Decision Making Processes1. https://doi.org/105465/255641. 2017;17: 605–621. doi:10.5465/255641

16. Dalkey N, Helmer O. An Experimental Application of the DELPHI Method to the Use of Experts. http://dx.doi.org/101287/mnsc93458. 1963;9: 458–467. doi:10.1287/MNSC.9.3.458

17. Mlecnik B, Bifulco C, Bindea G, Marliot F, Lugli A, Lee JJ, et al. Multicenter International Society for Immunotherapy of Cancer Study of the Consensus Immunoscore for the Prediction of Survival and Response to Chemotherapy in Stage III Colon Cancer. Journal of Clinical Oncology. 2020;38: 3638. doi:10.1200/JCO.19.03205

18. Using the Delphi method | IEEE Conference Publication | IEEE Xplore. [cited 7 Nov 2022]. Available: https://ieeexplore.ieee.org/abstract/document/6017716

19. Controlling the False Discovery Rate: A Practical and Powerful Approach to Multiple Testing on JSTOR. [cited 21 Oct 2022]. Available: https://www.jstor.org/stable/2346101

20. Wright SP. Adjusted P-Values for Simultaneous Inference. Biometrics. 1992;48: 1005. doi:10.2307/2532694

21. Hochberg Y. A Sharper Bonferroni Procedure for Multiple Tests of Significance. Biometrika. 1988;75: 800. doi:10.2307/2336325

22. Ellis P. The essential guide to effect sizes: Statistical power, meta-analysis, and the interpretation of research results. 2010. Available: https://books.google.com/books?hl=en&lr=&id=UUcgAwAAQBAJ&oi=fnd&pg=PR13&dq=The+Essential+Guide+to+Effect+Sizes+&ots=-d7gkrhpeO&sig=xjGU7RQ1tikVViYt6QlI7LdtbQg

23. Sullivan GM, Feinn R. Using Effect Size—or Why the P Value Is Not Enough. J Grad Med Educ. 2012;4: 279. doi:10.4300/JGME-D-12-00156.1

24. Sawilowsky SS. New Effect Size Rules of Thumb. Journal of Modern Applied Statistical Methods. 2009;8: 597–599. doi:10.22237/jmasm/1257035100

25. Dickinson ME, Flenniken AM, Ji X, Teboul L, Wong MD, White JK, et al. High-throughput discovery of novel developmental phenotypes. Nature. 2016;537: 508–514. doi:10.1038/nature19356

26. Bradley A, Anastassiadis K, Ayadi A, Battey JF, Bell C, Birling MC, et al. The mammalian gene function resource: The International Knockout Mouse Consortium. Mammalian Genome. 2012;23: 580–586. doi:10.1007/s00335-012-9422-2

27. Brown SDM, Moore MW. The International Mouse Phenotyping Consortium: Past and future perspectives on mouse phenotyping. Mammalian Genome. 2012;23: 632–640. doi:10.1007/s00335-012-9427-x

28. Hrabĕ de Angelis M, Nicholson G, Selloum M, White JK, Morgan H, Ramirez-Solis R, et al. Analysis of mammalian gene function through broad-based phenotypic screens across a consortium of mouse clinics. Nat Genet. 2015;47: 969–978. doi:10.1038/ng.3360

29. Bryant CD, Zhang NN, Sokoloff G, Fanselow MS, Ennes HS, Palmer AA, et al. Behavioral Differences among C57BL/6 Substrains: Implications for Transgenic and Knockout Studies. J Neurogenet. 2008;22: 315. doi:10.1080/01677060802357388

30. Haselimashhadi H, Mason JC, Mallon AM, Smedley D, Meehan TF, Parkinson H. OpenStats: A robust and scalable software package for reproducible analysis of high-throughput phenotypic data. In: PLoS ONE [Internet]. 2020 [cited 21 Jan 2021]. doi:10.1371/journal.pone.0242933

31. Gałecki A, Burzykowski T. Linear Mixed-Effects Model. Springer. 2013. doi:10.1007/978-1-4614-3900-4_13

32. Team RC-VRC, 2013 undefined. R: A language and environment for statistical computing. yumpu.com. [cited 18 Oct 2022]. Available: https://www.yumpu.com/en/document/view/6853895/r-a-language-and-environment-for-statistical-computing

33. Haseli Mashhadi H. Bioconductor - OpenStats. 2022 [cited 21 Jan 2021]. doi:10.18129/B9.bioc.OpenStats

34. Viechtbauer W. Conducting Meta-Analyses in R with the metafor Package. J Stat Softw. 2010;36: 1–48. doi:10.18637/JSS.V036.I03

35. Bierer BE, Crosas M, Pierce HH. Data Authorship as an Incentive to Data Sharing. New England Journal of Medicine. 2017;376: 1684–1687. doi:10.1056/NEJMSB1616595

36. International Consortium of Investigators for Fairness in Trial Data Sharing, Devereaux PJ, Guyatt G, Gerstein H, Connolly S, Yusuf S. Toward Fairness in Data Sharing. N Engl J Med. 2016;375: 405–7. doi:10.1056/NEJMp1605654

## References

1. Karp NA, Mason J, Beaudet AL, Benjamini Y, Bower L, Braun RE, et al. Prevalence of sexual dimorphism in mammalian phenotypic traits. Nat Commun. 2017;8: 15475. doi:10.1038/ncomms15475

